# Ongoing amphibian trade into the United States threatens salamander biodiversity

**DOI:** 10.1101/2023.01.05.522946

**Authors:** Patrick J. Connelly, Noam Ross, Oliver C. Stringham, Evan A. Eskew

## Abstract

The fungal pathogen *Batrachochytrium salamandrivorans* (*Bsal*) is a major potential threat to salamander biodiversity in North America, where it is not yet known to occur. In the United States, a 2016 policy restricted the trade in 20 salamander genera in attempts to prevent *Bsal* introduction. However, little comprehensive data is available to evaluate the impact of this policy action. Here, we collated a dataset of United States amphibian imports from 1999 to 2021 and show that reported legal trade in the targeted taxa was effectively reduced by the ban. Unfortunately, amphibian trade into the United States continues to risk *Bsal* introduction given that other species and genera now known to carry *Bsal* are still traded in large quantities (millions of live individuals annually). Additional policy responses focused on *Bsal* carrier taxa, especially frogs in the genus *Rana*, could help mitigate the impact of *Bsal* on North American salamanders.

## INTRODUCTION

The emerging infectious disease chytridiomycosis has devastated amphibian populations globally (Fisher and Garner 2020). The disease is caused by two closely-related fungal pathogens, *Batrachochytrium dendrobatidis* (*Bd*) and *Batrachochytrium salamandrivorans* (*Bsal*). First described in 1998 (Berger et al. 1998), *Bd* has been implicated in the decline of hundreds of amphibian species, including the presumed extinction of over 90 (Skerratt et al. 2007, Scheele et al. 2019; but see Lambert et al. 2020). *Bd* has a broad host range and is capable of infecting members of all three amphibian orders (Berger et al. 2016), but population-level impacts of *Bd* outbreaks have been mostly concentrated in frogs (order Anura) (Skerratt et al. 2007, Scheele et al. 2019). By contrast, *Bsal*, which was only recognized as a distinct species in 2013 (Martel et al. 2013), has caused significant declines in western European salamander populations. Notably, the fire salamander (*Salamandra salamandra*) has experienced mass die-offs and local extirpations from *Bsal* (Martel et al. 2013, Spitzen-van der Sluijs et al. 2013, Scheele et al. 2019). While *Bsal* was historically understood to exclusively infect salamanders (order Caudata) (Martel et al. 2014), we now know it is also capable of infecting anurans, suggesting that frogs could act as important vectors of the fungus (Nguyen et al. 2017, Stegen et al. 2017). *Bsal* is thought to be endemic to Asia, and while some Asian amphibians may therefore act as aclinical *Bsal* reservoir hosts due to long-term coexistence with the pathogen (Martel et al. 2014), movement of *Bsal* into novel geographic regions is expected to expose largely naïve amphibian populations that could become imperiled by disease.

The global wildlife trade is a primary mechanism for the international spread of *Batrachochytrium* species (Weldon et al. 2004, Fisher et al. 2009, Schloegel et al. 2009, Schloegel et al. 2012, Martel et al. 2014, O’Hanlon et al. 2018, Fu and Waldman 2022). *Bd* has been detected on amphibians in diverse trade endpoints, including pet stores, food markets, and scientific/zoological institutions, reflecting high demand for frogs in these settings (Daszak et al. 2003, Schloegel et al. 2009, Wombwell et al. 2016). Similarly, the international trade in salamanders sourced from Asia has been implicated as an important pathway for *Bsal* spread (Martel et al. 2014), and *Bsal* has been detected on animals in pet stores and captive collections in Europe (Martel et al. 2014, Sabino-Pinto et al. 2015, Nguyen et al. 2017, Fitzpatrick et al. 2018, Sabino-Pinto et al. 2018). *Bsal*’s high virulence and broad host range suggest there may be serious impacts on salamanders globally should it continue to spread via wildlife trade on a scale similar to *Bd* (Martel et al. 2014, Gray et al. 2015, Yap et al. 2015, Richgels et al. 2016).

Amphibian conservationists have been particularly worried about the consequences of *Bsal* introduction to the United States given the country’s salamander biodiversity. The United States hosts nearly 30% of all salamander species globally (Yap et al. 2015), and prospective laboratory studies on native North American amphibians indicate that numerous species are susceptible to lethal *Bsal* infection (Carter et al. 2020, Wilber et al. 2021, Gray et al. 2022). In light of its potentially large pool of *Bsal*-susceptible salamander species, the robust amphibian trade into the United States raises serious concerns. The live amphibian trade into the country numbers in the millions of individuals annually (Schloegel et al. 2009, Herrel and van der Meijden 2014, Altmann and Kolby 2017), and an analysis focused on potential *Bsal* carrier species estimated that ∼750,000 of these individuals entered the United States in the five-year period between 2010 and 2014 (Yap et al. 2015).

Thankfully, there has been policy action in the United States aimed at mitigating the threat of *Bsal* introduction. In 2016, the United States Fish and Wildlife Service (USFWS) published an interim rule listing 201 salamander species from 20 genera under the Lacey Act as “injurious wildlife,” effectively prohibiting their importation into the United States (US Fish and Wildlife Service 2016). Ostensibly, this policy has been a success: in the years since the Lacey Act action, survey efforts have not detected *Bsal* in captive (Klocke et al. 2017) or wild amphibians (Waddle et al. 2020, Hill et al. 2021) in the United States. However, this encouraging news is tempered by the fact that numerous amphibian genera not considered in the interim rule have since been found to be capable of hosting *Bsal*, including notable anuran genera such as *Alytes, Bombina, Hyla*, and *Rana* (Nguyen et al. 2017, Stegen et al. 2017, Schulz et al. 2020, Gray et al. 2022). As a consequence, the importation of potential *Bsal* carrier species into the United States likely continues, with little published data quantifying changes in the amphibian trade following the 2016 Lacey Act interim rule.

Here, we collate and clean a comprehensive dataset spanning two decades of amphibian imports into the United States to evaluate trends in amphibian trade, with a focus on species potentially driving the global spread of *Bsal*. More specifically, this multi-decade dataset allows us to address multiple questions of conservation interest: 1) How has the overall magnitude of amphibian imports into the United States changed over time? 2) From which countries do United States amphibian imports originate, and are these countries known to have *Bsal*? 3) Which United States ports do amphibian imports flow into? 4) Did the 2016 Lacey Act interim rule effectively restrict trade in the listed species? 5) What is the magnitude of trade in genera and species not currently listed as injurious in the United States that may nonetheless serve as *Bsal* carriers? Collectively, our results demonstrate how the global amphibian trade continues to generate opportunities for *Bsal* invasion into the United States.

## METHODS

To obtain a complete amphibian trade dataset for analysis, we cleaned and combined three smaller datasets that together gave a continuous record of amphibian imports to the United States from 1999 to 2021. All of these primary data sources contain data from the USFWS Law Enforcement Management Information System (LEMIS). LEMIS data have been widely used by researchers interested in various taxa to help understand wildlife trade that flows into or out of the United States (Smith et al. 2017, Marshall et al. 2020), and the dataset has been praised as a model wildlife trade surveillance system given the relative specificity of information contained therein (Hughes et al. 2021). Historically, however, researchers have obtained LEMIS data through repeated Freedom of Information Act (FOIA) requests, with different groups requesting different information on different taxa over different time periods. Thus, compilation of a complete LEMIS amphibian trade dataset required significant data collation efforts, as described here. Our first major dataset was a previously published collection of curated LEMIS data that covers the wildlife trade, broadly considered, from 2000 to 2014 (Eskew et al. 2020). We supplemented this general wildlife trade time series with a specific FOIA request to the USFWS for LEMIS amphibian import data from 2016 to 2021. We requested and received data that largely matched the LEMIS fields and formatting described in Eskew et al. 2020. Finally, we were able to add two missing years of data, 1999 and 2015, to our time series using LEMIS data that the USFWS has recently made publicly available (https://www.fws.gov/library/collections/office-law-enforcement-importexport-data). We combined these three primary data sources by reconciling field names in all datasets to match the Eskew et al. 2020 data. We also filtered the general wildlife trade datasets to only represent amphibian trade data. For the Eskew et al. 2020 data source, the “taxa” field allows for easy subsetting to rows with “amphibian” values. For the 1999 and 2015 LEMIS data, we used the wildlife category field to recover any rows with “Amp” or “AMP” values, indicating amphibian records. However, some rows in this dataset have no assigned wildlife category, and we didn’t want to overlook potential amphibian records simply because they were not labeled as such. Therefore, we used the current amphibian taxonomy available from AmphibiaWeb (https://amphibiaweb.org/taxonomy/AWtaxonomy.html) to recover any records that were missing wildlife category data but whose listed genus matched with a known amphibian genus. During this process, we matched on currently accepted AmphibiaWeb genera and any known generic synonyms. Finally, we cleaned all datasets following the basic protocol outlined in Eskew et al. 2020 (i.e., ensured values matched with valid codes provided by the USFWS). In summary, these data cleaning steps left us with three cleaned amphibian trade datasets containing a consistent set of variables and field values.

After combining our three datasets to obtain a complete time series of amphibian imports from 1999 to 2021, we harmonized the taxonomic information contained within these records. The taxonomic information in LEMIS is subject to data entry errors but may also represent shifts in preferred nomenclature over time (Eskew et al. 2020). For example, in our full amphibian trade dataset, the American bullfrog initially appeared with the scientific name *Rana catesbeiana*, the misspelling *Rana catesbeinana*, and the synonym *Lithobates catesbeianus*. To conduct the most accurate, complete analyses, we sought to reconcile this sort of competing nomenclature. As such, we cleaned all of the taxonomic names in our dataset, using the AmphibiaWeb taxonomy as our reference given that AmphibiaWeb is one of the most widely-used resources by amphibian biologists (Womack et al. 2022). We first used the AmphibiaWeb taxonomy to automatically convert known synonyms to the preferred AmphibiaWeb nomenclature, where possible. At that point, comparison of the scientific names in our trade dataset to the AmphibiaWeb taxonomy alerted us to potential instances of taxonomic misspellings, which we manually corrected. Once we synonymized and cleaned all amphibian names, we were then able to add higher-level taxonomic information to the data, namely taxonomic family and order.

To summarize, our collated amphibian trade dataset consists of 90,641 records representing over 43,000 wildlife shipments from 1999 to 2021 (Connelly et al. 2023). While prior work has sometimes provided longer time series of United States amphibian trade (Olsen et al. 2019), these efforts have relied on CITES data which fail to capture trade in the many species that are not listed on the CITES Appendices (Eskew et al. 2019, Hughes et al. 2021). By contrast, the comprehensive LEMIS dataset more fully captures trends in United States amphibian imports. As a result of our data cleaning procedures, 74.9% of our import records were assigned complete scientific names recognized by AmphibiaWeb (note that 24.3% of records are only reported to the genus level [“sp.” for the species field] and therefore cannot have complete scientific names). These scientific name matches represent 935 unique amphibian species. Over 94% of records were either originally assigned a generic name or had a cleaned generic name that matched with AmphibiaWeb genera.

Our analyses also required us to gather data on the amphibian species listed under the 2016 Lacey Act interim rule, those more broadly that could serve as potential *Bsal* carriers, and the countries in which *Bsal* is known to exist in wild amphibians. A list of amphibian species and genera listed under the Lacey Act interim rule was pulled directly from publicly available information from USFWS (US Fish and Wildlife Service 2016). To gather data on *Bsal* carrier species (Supporting Information), we conducted a review of the primary and gray literature, building off previous work (Grear et al. 2021). In reviewing the *Bsal* literature, we noted the amphibian species known to harbor *Bsal* infections, whether or not disease symptoms were also observed in those species, and the context in which the infection was detected (i.e., in a wild, captive, or laboratory-exposed animal). In our analyses, we were maximally conservative and considered a potential *Bsal* carrier species to be any amphibian species with a documented *Bsal* infection, irrespective of disease symptoms. Additionally, during the course of our literature review, we recorded the countries in which *Bsal* was detected in wild amphibians. Here, the status of Hong Kong deserves special note. LEMIS amphibian trade data record Hong Kong as a distinct country of origin despite the fact that Hong Kong is technically a special administrative region of China. Although we are unaware of *Bsal* detections in wild amphibians from Hong Kong specifically, *Bsal* is known from the abutting Chinese province of Guangdong (Yuan et al. 2018). As such, we consider Hong Kong as *Bsal*-positive for all relevant analyses, but report its trade exports as distinct from that of China generally, which is consistent with the presentation in the LEMIS data. Our decision here is further justified by the fact that Hong Kong is known to be among the most important amphibian exporters to the United States and transports animals infected with *Bd* (Kolby et al. 2014, Sinclair et al. 2021). Thus, lumping Hong Kong together with China could obscure its unique role in the amphibian trade.

For analysis of trends in amphibian trade over time, we focused primarily on trade involving live animals (records coded as “LIV” in the “description” field), given these are the wildlife shipments that are arguably the most likely to transport *Bsal* internationally (Gratwicke et al. 2010, Altmann and Kolby 2017). We visualized trends in live amphibian imports over time, highlighting major divisions in the data, including order of the amphibians, country of origin, and port of entry. In particular, we sought to highlight trade patterns in the post-Lacey Act period from 2017 to 2021 (although the interim Lacey Act rule was put into effect in 2016, 2017 was the first full year in which United States trade was subject to the restrictions). Next, we examined trade in the amphibian taxa included in the Lacey Act interim rule. Here, we chose to analyze trade data at the level of genera rather than species to be conservative: because a relatively large proportion of LEMIS records in our final dataset (24.3%) are only reported with the ambiguous species designation of “sp.”, analyses at the genus level may be necessary to fully capture relevant trade patterns in key Lacey Act taxa. Finally, we highlighted trade in potential *Bsal* carrier species and the genera they belong to. Where relevant, we statistically analyzed temporal trends in amphibian trade with linear models implemented within R version 4.1.1 (R Core Team 2021), using yearly counts of imported live individuals as the response variable and year as a continuous explanatory variable. All data visualizations were created using ‘ggplot2’ (Wickham 2010) and ‘cowplot’ (Wilke 2020).

## RESULTS

From 1999 to 2021, the live amphibian trade into the United States totaled 85.82 million individuals, an average of 3.73 million individuals annually (Figure 1). 31.28 million of these live amphibian individuals (36.4%) were declared as originating from the wild. Despite the fact that the live amphibian trade has declined significantly during this timeframe (R^2^ = 0.58, *F*(1, 21) = 31.41, *p* < 0.001), in the post-Lacey Act period (i.e., 2017-2021) there were still, on average, 2.84 million live amphibians imported annually (Figure 1). The trade in live amphibians is accompanied by a robust trade in amphibian legs and meat: our dataset indicates 2.76 million kg of amphibian legs/meat entered the United States annually from 1999 to 2021. Over 90% of the total amphibian leg and meat trade in this time period (58.17 million kg) derived solely from one species, the American bullfrog (*R. catesbeiana*).

**Figure 1.**
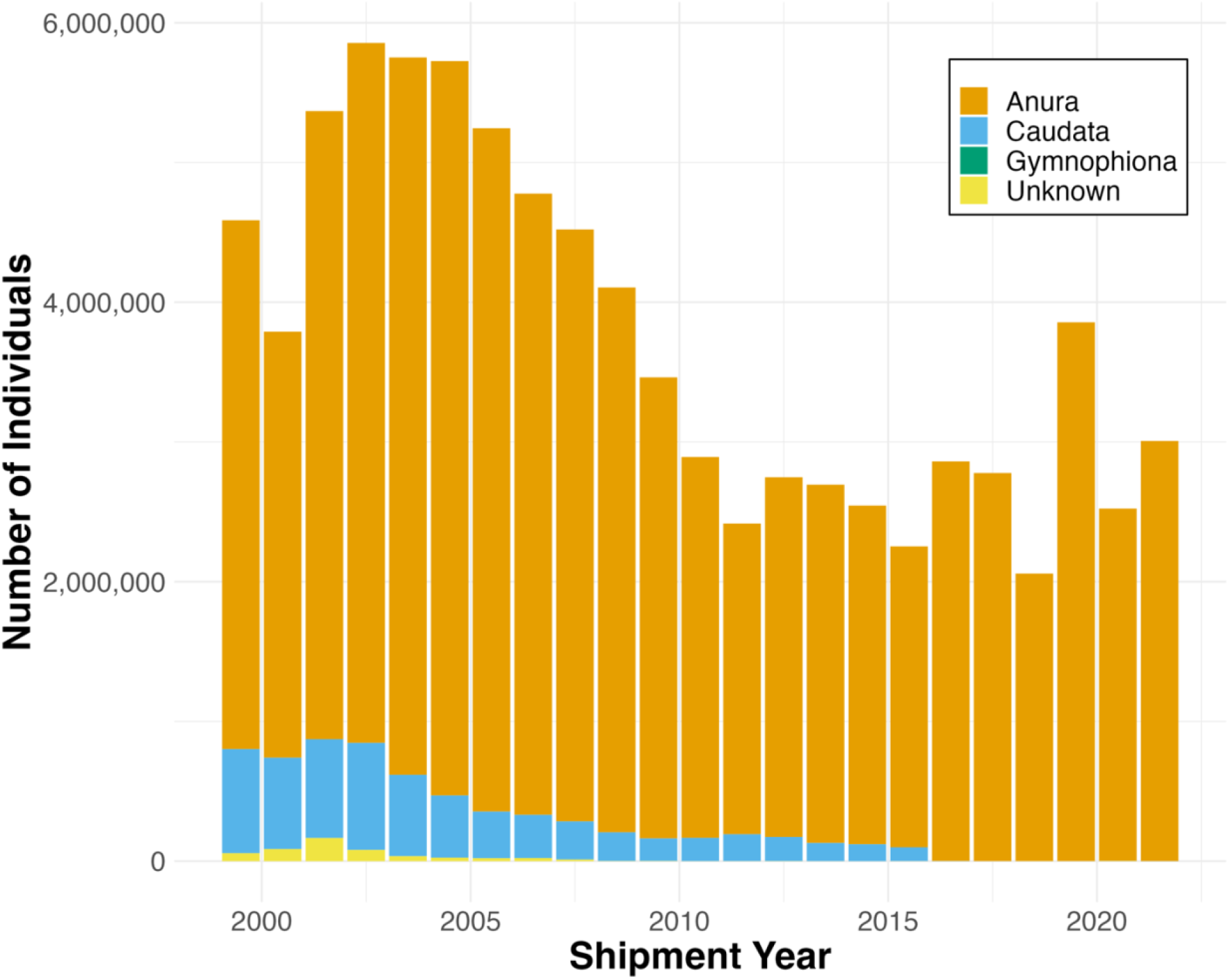
Live amphibian imports to the United States from 1999 to 2021 by taxonomic order. This figure shows only LEMIS import data for live amphibian shipments that were recorded in terms of numbers of individuals. Colors indicate the amphibian taxonomic order, and the ordering of categories within each stacked bar corresponds to ordering in the legend. Note that caecilians (order Gymnophiona) represent an extremely small portion (<1%) of live amphibian imports annually.

The live amphibian trade into the United States is dominated by members of the order Anura. Anurans represented >80% of the live imported amphibians in every year studied, rising to >99% of individuals in the post-Lacey Act period (Figure 1). Caudates always represented <20% of individuals imported annually, with this number dropping to <1% in the years from 2017 to 2021 (Figure 1). Members of Gymnophiona represented <1% of live amphibian individuals traded in every year studied (Figure 1).

Amphibians imported to the United States were primarily exported from countries in Asia and Latin America. Nearly one-half (48.5%) of all live amphibians imported to the United States from 1999 to 2021 originated from just two Asian exporters: Taiwan (30.04 million individuals; 35.0%) and Hong Kong (11.57 million individuals; 13.5%). Ecuador, Singapore, and China round out the top five amphibian exporters to the United States during this time period. Further, >50% of all live amphibian individuals imported annually into the United States originated from countries that are known to host wild amphibians infected with *Bsal*, with the exception of the years 1999, 2011, and 2012 (Figure 2). In 2017 and all subsequent years, >64% of all live imported amphibian individuals have originated from *Bsal*-positive countries (Figure 2). Taiwan, as a *Bsal*-positive country, emerges as a particularly important amphibian exporter to the United States: in the post-Lacey Act period, Taiwan is the point of origin for >58% of live amphibians imported each year (Figure 2). We further note that if we consider all wildlife imports to the United States from 2000 to 2014 (using the full Eskew et al. 2020 dataset), Taiwan is only responsible for the export of ∼4.3% of live individuals. Thus, Taiwan appears to play a disproportionate role in the amphibian trade specifically.

**Figure 2.**
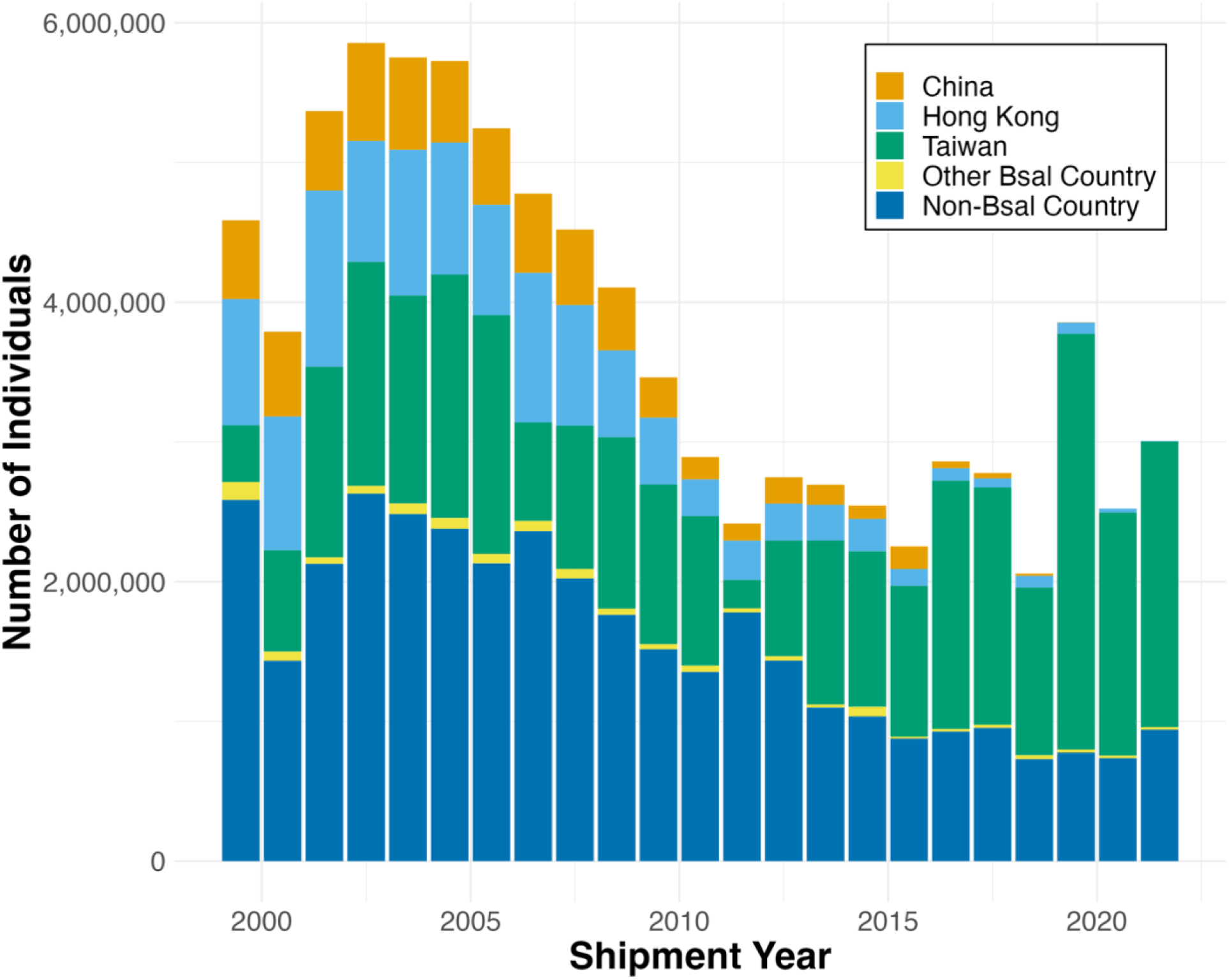
Live amphibian imports to the United States from 1999 to 2021 by country of origin. This figure shows only LEMIS import data for live amphibian shipments that were recorded in terms of numbers of individuals. Colors indicate the country of origin, and the ordering of categories within each stacked bar corresponds to ordering in the legend. We specifically highlight the 10 countries that are known to host wild, *Bsal*-positive amphibians, but for simplicity of presentation, seven of these countries (Belgium, Germany, Japan, Netherlands, Spain, Thailand, and Vietnam) have been combined into an “Other Bsal Country” category.

Although live amphibians imported to the United States flow into over 40 unique ports of entry, the majority concentrate in relatively few major ports nationwide. From 1999 to 2021, the top five United States ports of entry for live amphibians have been Los Angeles (39.11 million individuals; 45.6%), New York (17.97 million individuals; 20.9%), San Francisco (12.88 million individuals; 15.0%), Brownsville (4.34 million individuals; 5.1%), and Miami (2.51 million individuals; 2.9%). Since the year 2000, these five ports alone have accounted for >80% of all live amphibians imported annually into the United States (Figure 3). In the post-Lacey Act period, Los Angeles has been the port of entry for >68% of all live amphibians imported annually (Figure 3). These amphibian import patterns largely mirror the more general wildlife trade. Over the 2000 to 2014 period for which we have comprehensive data on the overall wildlife trade, Los Angeles was the port of entry for nearly 50% of live individuals, while New York and Miami accounted for ∼14% and ∼13% of individuals, respectively.

**Figure 3.**
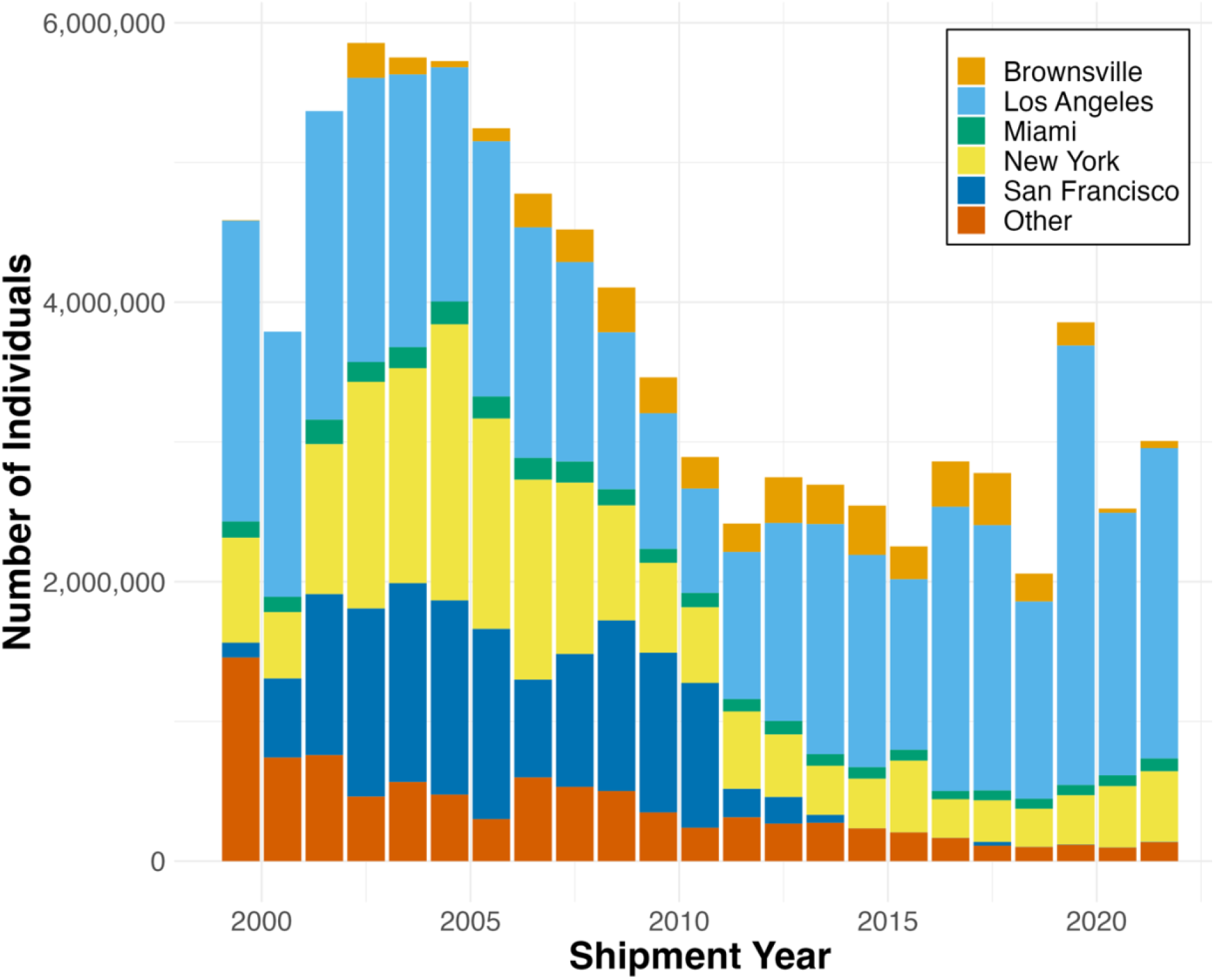
Live amphibian imports to the United States from 1999 to 2021 by port of entry. This figure shows only LEMIS import data for live amphibian shipments that were recorded in terms of numbers of individuals. Colors indicate the United States port of entry, highlighting the five most common ports of entry for live amphibians, and the ordering of categories within each stacked bar corresponds to ordering in the legend.

Of the 20 caudate genera listed under the 2016 Lacey Act interim rule, 17 were shipped live into the United States at some point during the study period (Figure 4a). Import of these genera averaged 255,011 live individuals per year, but there was a strong decrease in trade over time even before the Lacey Act interim rule was announced (model estimated decrease of 34,821 individuals per year; R^2^ = 0.87, *F*(1, 21) = 145.6, *p* < 0.001). Fewer than 200 individuals from these genera were imported annually in the post-Lacey Act period (Figure 4b). Moreover, the metadata regarding these shipments indicates that the vast majority were treated as expected given the Lacey Act interim rule: the only shipments containing Lacey Act genera that were actually cleared for entry to the United States (as opposed to refused) were declared as being traded for scientific purposes with the exception of a single *Cynops pyrrhogaster* individual that was traded for “personal” purposes and was reported as cleared for entry in 2017.

**Figure 4.**
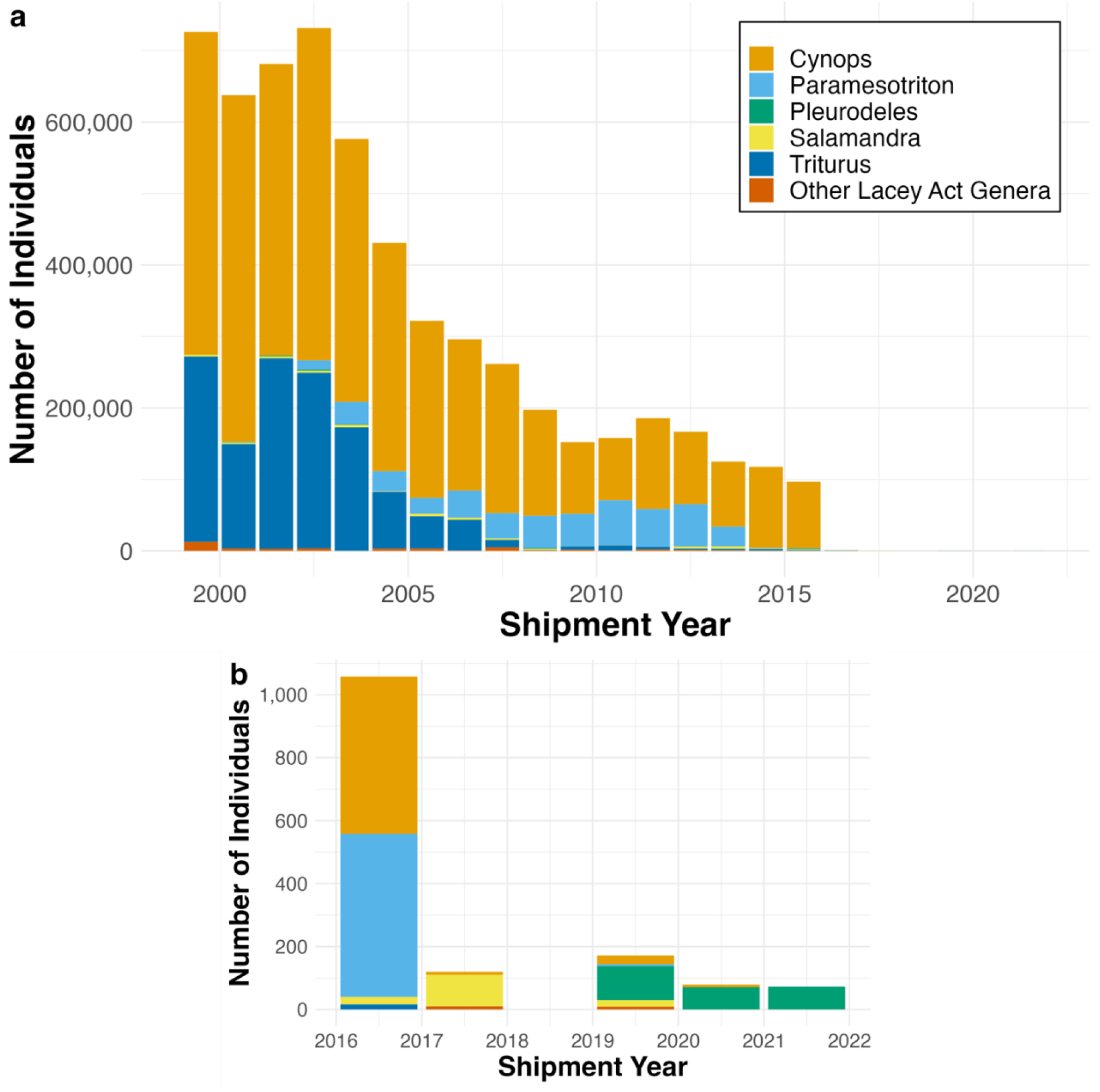
Live amphibian imports to the United States from 1999 to 2021 of genera listed under the 2016 Lacey Act interim rule. This figure shows only LEMIS import data for live amphibian shipments that were recorded in terms of numbers of individuals. Data are shown for both (a) the full 1999-2021 timeline and (b) the 2016-2021 period corresponding to the year of the Lacey Act interim rule and subsequent years (note difference in y-axis scales). Colors indicate the amphibian genera, and the ordering of categories within each stacked bar corresponds to ordering in the legend.

Our review of the literature identified 85 potential *Bsal* carrier species spanning 43 genera and two amphibian orders (Anura and Caudata; Supporting Information). We found a modest trade in these specific species in the post-Lacey Act period. Throughout the 1999 to 2021 time period, 29 potential *Bsal* carrier species have been imported to the United States, but the data indicate that fewer than 150 live individuals of these species have been traded annually since 2016 (Figure 5). However, examination of live trade in the genera that potentially carry *Bsal* paints a vastly different picture. Imports of these genera have averaged 2.44 million live individuals annually in the post-Lacey Act period (Figure 6a). Put differently, more than 80% of the United States’ live amphibian imports annually (represented in full in Figure 1) derives from amphibian genera that have members known to host *Bsal*. This trade is dominated by the genus *Rana*, which represents >97% of the live trade in *Bsal* carrier genera every year from 2017 to 2021 (Figure 6a). The only other genera to represent more than 1% of annual trade over this time period was the genus *Bombina* in the years 2017 and 2018 (Figure 6a). In the post-Lacey Act period, over 70% of these individuals annually have their point of origin in Taiwan (Figure 6b), and over 70% annually are imported to the United States via the port of Los Angeles (Figure 6c).

**Figure 5.**
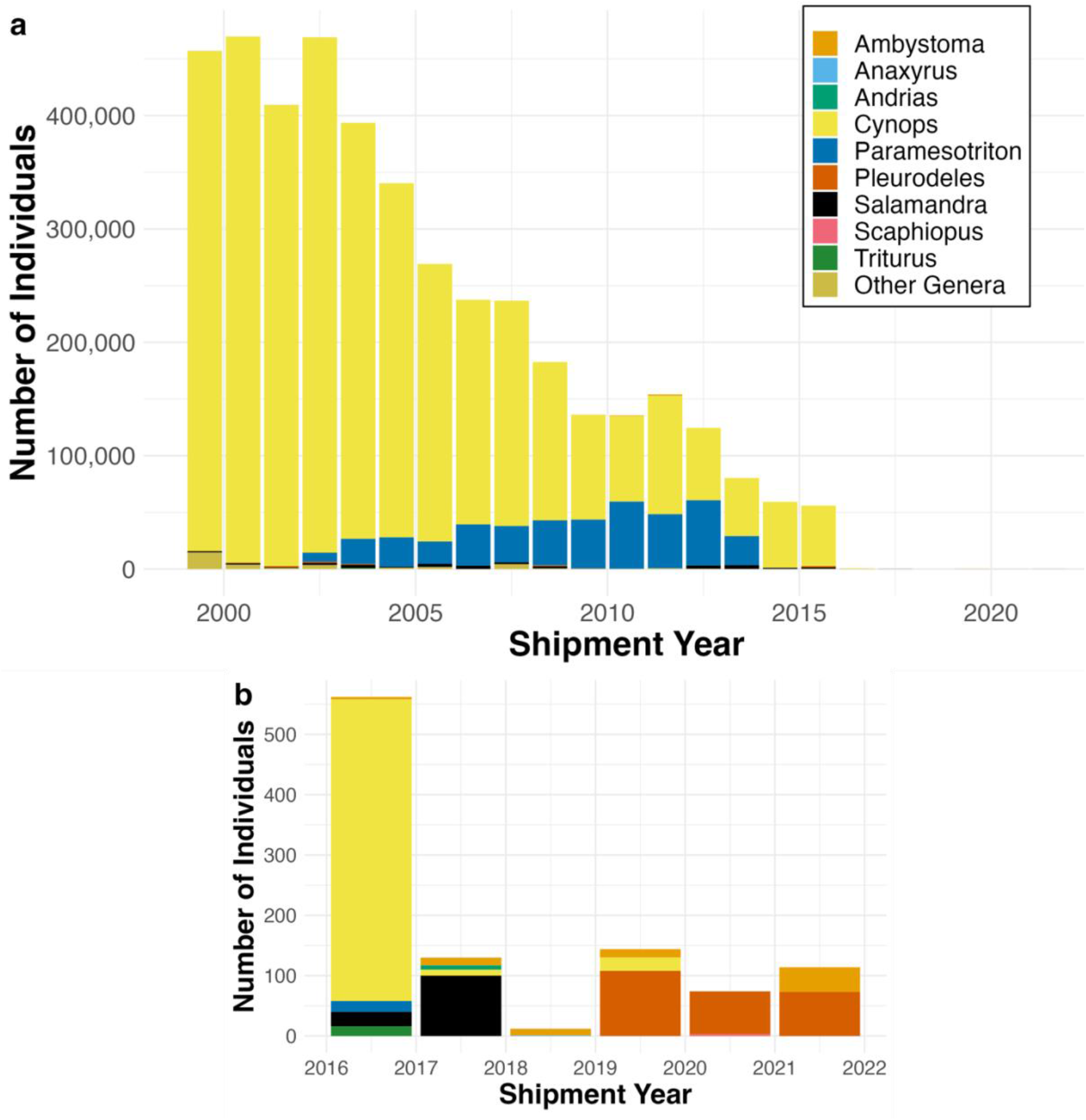
Live amphibian imports to the United States from 1999 to 2021 of known *Bsal* carrier species. This figure shows only LEMIS import data for live amphibian shipments that were recorded in terms of numbers of individuals. The LEMIS data were filtered to only those species we recorded as known *Bsal* carriers (n = 29 species in the LEMIS dataset). Data are shown for both (a) the full 1999-2021 timeline and (b) the 2016-2021 period corresponding to the year of the Lacey Act interim rule and subsequent years (note difference in y-axis scales). For visual simplicity, colors indicate the amphibian genera to which these species belong, and the ordering of categories within each stacked bar corresponds to ordering in the legend.

**Figure 6.**
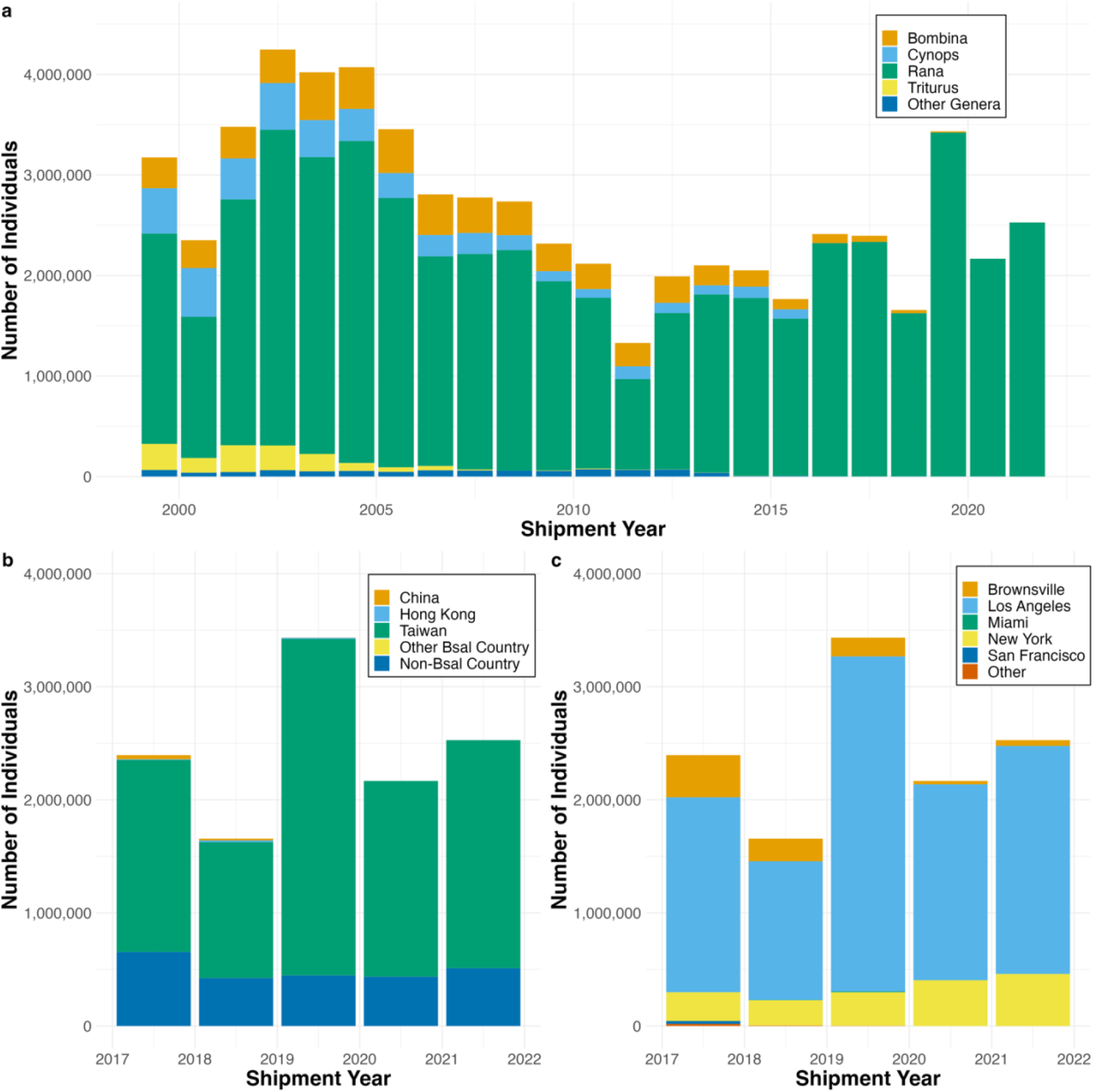
Live amphibian imports to the United States from 1999 to 2021 of known *Bsal* carrier genera. This figure shows only LEMIS import data for live amphibian shipments that were recorded in terms of numbers of individuals. The LEMIS data were filtered to only those genera we recorded as containing known *Bsal* carrier species (n = 32 genera in the LEMIS dataset). In (a) the full 1999-2021 timeline is shown. Colors in (a) indicate the amphibian genera, and the ordering of categories within each stacked bar corresponds to ordering in the legend. Panels (b) and (c) depict the 2017-2021 period corresponding to the years following the Lacey Act interim rule (note difference in y-axis scales), with (b) highlighting the country of origin for *Bsal* carrier genera and (c) highlighting the United States port of entry for *Bsal* carrier genera.

## DISCUSSION

Here, we collated and analyzed what is, to our knowledge, the longest comprehensive dataset of amphibian trade flows into the United States, and we evaluate this trade with respect to the potential for introduction of the amphibian pathogen *Bsal*. Our results show that while conservation policy has effectively reduced the trade in some known *Bsal* carrier species, other potential *Bsal* carrier taxa are still imported in large numbers, representing a continued threat to salamander biodiversity in the United States and North America more broadly (Gray et al. 2015, Yap et al. 2015, Richgels et al. 2016, Grear et al. 2021). These specific, conservation-relevant findings are embedded within the broader context of the overall United States amphibian trade that our dataset documents, and having this curated resource openly available will facilitate future trade-related research.

Generally, our analyses illustrate that the live amphibian trade into the United States is substantial and dominated by anurans. Despite there being a significant negative trend in the number of live amphibians entering the United States between 1999 and 2021, the most recent five years of data indicate that nearly three million live amphibians still enter the country annually, the vast majority (>99%) of them being anurans. These results align with other work suggesting the relative dominance of the anuran trade both in the United States and globally (Grear et al. 2021, Hughes et al. 2021). The prevalence of anurans in the amphibian trade along with their potential to carry *Bsal* requires us to reconsider the true magnitude of *Bsal* importation risk to the United States, as we discuss in more detail below.

The data also reveal that relatively few ports of entry concentrate amphibian flows entering the United States, an observation that could help guide effective allocation of disease surveillance and biosecurity effort (Sinclair et al. 2021). In particular, Los Angeles and New York represent the two most common ports for live amphibians to enter the United States in the post-Lacey Act period. As others have emphasized (Yap et al. 2015), Los Angeles is especially important as a potential entry point for *Bsal* to the United States: the majority of live individuals belonging to *Bsal* carrier genera that entered the country in the post-Lacey Act period did so through this port. Los Angeles County also has a very high number of pet-related businesses that could act as dispersal hubs, distributing infected animals further across the landscape (Richgels et al. 2016, Sinclair et al. 2021).

In addition, the LEMIS data provide information on the country of origin of amphibian imports to the United States. Other work has suggested that United States amphibian imports are becoming increasingly spatially consolidated, with imported animals coming from fewer exporting countries over time (Sinclair et al. 2021). Our data are consistent with this observation given that a single exporting country, Taiwan, has sourced the majority of live amphibian imports each year since 2016. This finding is particularly troubling given that wild Taiwanese amphibians are known to be infected with *Bsal* (Beukema et al. 2018). However, interpretation of these import patterns requires some caution given that the country of origin listed in the LEMIS data may, in some cases, erroneously report a country of reexport rather than the shipment’s true country of origin (Sinclair et al. 2021). Nonetheless, like information on common ports of entry, the best available data on country of origin for amphibian imports could be used to prioritize disease surveillance efforts by allowing investigators to target imported amphibian shipments from high-risk *Bsal* areas for pathogen screening. Alternatively, there may be opportunities for productive bilateral cooperation on more comprehensive programs that screen amphibians for *Bsal* at both the country of export and the United States port of entry.

A primary aim of our analyses was to evaluate the impact of the 2016 Lacey Act interim rule on trade in the species listed therein. We found that the Lacey Act action appears to have been highly effective in reducing trade in the targeted species (Grear et al. 2021). Since 2017, the LEMIS data suggests that only a few hundred of these individuals have entered the country in total, nearly all intended for scientific purposes. Given the frequent failure of the conservation community in translating research and scientific understanding to real-world solutions (Grant et al. 2019), the 2016 Lacey Act interim rule should be commended as a tangible conservation success. The interim rule was implemented quickly relative to the discovery of the *Bsal* threat (Altmann and Kolby 2017), and the scope of the action was far-reaching, given information available at the time. The apparent efficacy of the Lacey Act interim rule is tempered by a few caveats, however. First, in other wildlife trade contexts, shipment declarations and subsequent data sources may contain purposeful misreporting that conceals the true geographic origin, source (i.e., captive-bred vs. wild-caught), or even species of the shipment in question (Nijman and Shepherd 2011, Altmann and Kolby 2017, Smith et al. 2017, Harfoot et al. 2018, Hierink et al. 2020), perhaps especially when such misreporting helps the shipment to evade regulation. As such, there is no way to guarantee that injurious species listed in the interim rule have not been imported by declaring fraudulent taxonomic information. Second, our data does not account for the potential illegal trade in Lacey Act listed species. The illegal wildlife trade is inherently difficult to characterize (Rosen and Smith 2010, Fukushima et al. 2021), but it remains a possible avenue for *Bsal* introduction to the United States despite the apparent success of Lacey Act enforcement. In fact, we might expect illegal trade frequency to increase for listed *Bsal* carrier species given that trade regulations (such as trade bans) can act to inflate prices and drive trade underground in cases where consumer demand remains high (Challender and MacMillan 2014).

Setting aside potential problems with comprehensive enforcement of the 2016 Lacey Act interim rule, the most significant issue with the current regulation is simply that the rule was written with incomplete information on *Bsal* host range. As a result, import of potentially injurious taxa to the United States continues. Of particular concern is continued trade of six anuran genera now known to carry *Bsal*: *Alytes, Anaxyrus, Bombina, Hyla, Rana*, and *Scaphiopus*. While previous analyses have highlighted the potential *Bsal* introduction risk associated with *Bombina*, a *Bsal* carrier genus popular in the pet trade and native to Eurasia (Grear et al. 2021, Fu and Waldman 2022), they have not fully addressed issues related to the genus *Rana*. The American bullfrog (*R. catesbeiana)* is one of the most heavily-traded species within the genus and is thought to have played a major role in the global spread of *Bd* (Schloegel et al. 2009, Schloegel et al. 2012, O’Hanlon et al. 2018, Yap et al. 2018, Byrne et al. 2019). Preliminary data indicates the American bullfrog may be resistant to *Bsal* (Gray et al. 2022), but other members of *Rana* appear to be capable of carrying the pathogen in laboratory (Gray et al. 2022) and field settings (Schulz et al. 2020) (although we note that others have interpreted some of these findings with caution [Grear et al. 2021]). In sum, the current evidence base, which has been generated by independent research groups, suggests that at least some members of *Rana* are susceptible to *Bsal*, and further investigations are urgently needed to more comprehensively assess the *Bsal* carrier status of various *Rana* species, including the American bullfrog. If any *Rana* species commonly harbor *Bsal* this would be a critical concern for the amphibian conservation community given the broad distribution of many of these frogs and their popularity in the wildlife trade.

A multi-pronged approach may be used to help mitigate the threat of *Bsal* introduction to and spread within the United States. First, although large-scale surveillance efforts have thus far failed to detect *Bsal* in wild amphibians in the United States (Waddle et al. 2020, Hill et al. 2021), ongoing disease surveillance in multiple settings (i.e., in the field and along multiple nodes of the amphibian trade network) is imperative to ensure early detection and potential control of the pathogen is possible (Gray et al. 2015, Grear et al. 2021). Surveillance efforts may be most feasible and effective if they are targeted towards known *Bsal* carrier species, and, in the wildlife trade context, if they focus on major ports for amphibian entry (i.e., Los Angeles) and shipments coming from countries with a robust amphibian trade and known *Bsal* endemism (i.e., Taiwan). Despite the considerable logistical, economic, and political challenges, more widespread disease surveillance at ports of entry generally would boost the likelihood of detecting invasive pathogens before they are able to spread among native biota (Altmann and Kolby 2017, Fu and Waldman 2022). Further, this strategy is likely more cost-effective than widespread field surveillance efforts to detect *Bsal*, and it could be made even more comprehensive if source countries also engage in disease testing at the point of origin. Second, in addition to increased disease surveillance, amphibians within the United States stand to benefit from additional policy responses that could reduce the probability of *Bsal* introduction, given that prevention of introduction is arguably the most effective lever available for reducing ultimate disease impact (Gray et al. 2015, Garner et al. 2016). Most obviously, current scientific knowledge regarding *Bsal*’s host range could be used to update the 2016 Lacey Act interim rule (Grear et al. 2021, Fu and Waldman 2022), supporting the intended purpose of that initial listing action. It is now clear that the regulations governing amphibian trade in the United States are based on incomplete information and do not account for the full set of species that should be considered injurious as a consequence of their *Bsal* host status. Any of the *Bsal* carrier species or genera recovered in our literature search are potential candidates for listing. However, the genus *Rana* is clearly of most immediate relevance given the substantial volume of trade in that genus, especially if additional *Rana* species are found to be capable of carrying *Bsal*. Stronger trade restrictions might even adopt a precautionary principle regarding amphibian import to the United States. For example, in the amphibian trade context more generally, where trade itself can lead to overexploitation and species endangerment, there have been calls for policies built on the premise that all species should be granted trade protections until it can be demonstrated that trade is sustainable (Hughes et al. 2021). Translated to the wildlife disease setting, this might look like a trade policy that preemptively restricts trade until it can be shown that an amphibian species is unlikely to serve as a host for *Bsal*.

In addition to efforts to prevent the future introduction of *Bsal*-infected individuals to the United States, captive animals that are already present in the country and could possibly harbor *Bsal* deserve our attention (Grear et al. 2021). Other notable fungal pathogens of wildlife may have been introduced to North America via captive animals (Ladner et al. 2022), and *Bsal* has been widely detected in amphibian collections in Europe where release of former pets has likely been a pathway for disease spread to wild animals (Martel et al. 2014, Fitzpatrick et al. 2018, Sabino-Pinto et al. 2018). Our data indicate that large numbers of individuals from *Bsal* carrier genera have been imported to the United States from 1999-2021 for commercial purposes (suggesting these animals likely entered the pet trade), including 5.33 million *Bombina*, 4.03 million *Cynops*, and 1.30 million *Triturus*. While effective import regulations are invaluable, as previously discussed, we cannot ignore this stock of captive animals already present in the United States that potentially contains *Bsal*-infected individuals. Considerable efforts should be made to engage with and educate the public and amphibian hobbyists, informing them of resources available for the ethical and safe surrender of unwanted pet amphibians. Further, it is clear that transport of *Bsal*-infected captive amphibians has facilitated *Bsal* invasion of Europe (Nguyen et al. 2017). Interestingly, while the 2016 Lacey Act interim rule initially banned interstate movement of listed species, this restriction was overturned by the United States Court of Appeals for the District of Columbia in a 2017 decision. Amending the Lacey Act to strengthen interstate travel restrictions on listed species could help minimize potential spread of *Bsal* among captive collections in the United States and/or between captive collections and the wild, but such a policy may find opposition among pet breeders and owners. Similarly, implementation of disease-related standards for the rearing and maintenance of captive amphibians could help ensure that only *Bsal*-free animals are exchanged between breeders, amphibian resellers, and pet owners.

Amphibians are the most threatened group of vertebrates on the planet, with over one-third of species at risk of extinction due to multiple threats, including habitat destruction, pollution, and infectious disease (Stuart et al. 2004, Wake and Vredenburg 2008, Grant et al. 2016). Lessons learned from the global spread of *Bd* and *Bsal’s* invasion of Europe suggest that amphibian conservation in the United States is at a critical juncture. We have the ability to proactively reduce the likelihood of, and simultaneously prepare for, a new amphibian biodiversity crisis in North America driven by disease (Gray et al. 2015, Yap et al. 2015, Richgels et al. 2016, Grear et al. 2021). To avert additional devastating impacts on amphibians, it is imperative that further conservation actions are taken and creative regulations are implemented to help mitigate the spread of *Bsal* within the United States.

## Acknowledgements

We thank the United States Fish and Wildlife Service and its numerous employees whose professional service has helped make the LEMIS trade data more widely available to the scientific community. In addition, we acknowledge the current and former affiliates of EcoHealth Alliance who contributed to data acquisition and cleaning for some of the publicly available data used here, including Peter Daszak, William B. Karesh, Jon Paul Rodríguez, Katherine F. Smith, Kristine M. Smith, Allison M. White, and Carlos Zambrana-Torrelio.

## Data Availability Statement

Raw and cleaned data as well as analysis scripts for this project are publicly available at https://github.com/ecohealthalliance/amphibian_trade. Additionally, these same materials have been archived on Zenodo (Connelly et al. 2023).

## Author Contributions

P.J.C. and E.A.E. conceived the idea for the paper. All authors made key contributions to data collection and curation. P.J.C. and E.A.E. analyzed the data and drafted the manuscript. All authors made comments and edits that shaped the final version of the paper.

